# Differentiation between Descending Thoracic Aortic Diseases using Machine Learning and Plasma Proteomic Signatures

**DOI:** 10.1101/2023.04.26.538468

**Authors:** Amanda Momenzadeh, Simion Kreimer, Dongchuan Guo, Matthew Ayres, Daniel Berman, Kuang-Yuh Chyu, Prediman K Shah, Dianna Milewicz, Ali Azizzadeh, Jesse G. Meyer, Sarah Parker

## Abstract

**Background:** Descending thoracic aortic aneurysms and dissections can go undetected until severe and catastrophic, and few clinical indices exist to screen for aneurysms or predict risk of dissection.

**Methods:** This study generated a plasma proteomic dataset from 75 patients with descending type B dissection (Type B) and 62 patients with descending thoracic aortic aneurysm (DTAA). Standard statistical approaches were compared to supervised machine learning (ML) algorithms to distinguish Type B from DTAA cases. Quantitatively similar proteins were clustered based on linkage distance from hierarchical clustering and ML models were trained with uncorrelated protein lists across various linkage distances with hyperparameter optimization using 5-fold cross validation. Permutation importance (PI) was used for ranking the most important predictor proteins of ML classification between disease states and the proteins among the top 10 PI protein groups were submitted for pathway analysis.

**Results:** Of the 1,549 peptides and 198 proteins used in this study, no peptides and only one protein, hemopexin (HPX), were significantly different at an adjusted p-value <0.01 between Type B and DTAA cases. The highest performing model on the training set (Support Vector Classifier) and its corresponding linkage distance (0.5) were used for evaluation of the test set, yielding a precision-recall area under the curve of 0.7 to classify between Type B from DTAA cases. The five proteins with the highest PI scores were immunoglobulin heavy variable 6-1 (IGHV6-1), lecithin-cholesterol acyltransferase (LCAT), coagulation factor 12 (F12), HPX, and immunoglobulin heavy variable 4-4 (IGHV4-4). All proteins from the top 10 most important correlated groups generated the following significantly enriched pathways in the plasma of Type B versus DTAA patients: complement activation, humoral immune response, and blood coagulation.

**Conclusions:** We conclude that ML may be useful in differentiating the plasma proteome of highly similar disease states that would otherwise not be distinguishable using statistics, and, in such cases, ML may enable prioritizing important proteins for model prediction.

## Introduction

Thoracic aortic aneurysms arise due to dysregulated growth and remodeling of the aorta in the segment spanning from the aortic root to the diaphragm^1^, which predispose the vessel wall to dissection and rupture. Aortic dissections occur when there is a loss of integrity, also described as a tear, in the intimal layer of the blood vessel. This tearing results in the formation of a ‘false lumen’ in the medial layer of the vessel, allowing aberrant blood flow patterns, risk of aneurysm formation/rupture, risk of thrombosis, and reduced perfusion of downstream tissues, all of which are associated with substantial morbidity and mortality^2^. The incidence of thoracic aortic aneurysms and thoracic aortic aneurysm dissections (TAAD) has risen over the past several decades, with an approximate doubling of new events between 1982^3^ and 2006^4^ (from 5.9 to 10.1 events per 100,000 individuals, respectively). Increased incidence may be attributed to a combination of improved diagnosis as well as increased prevalence of risk-factors such as atherosclerosis, hypertension, and an aging population^4^. TAADs are further categorized by anatomical region into ascending aneurysms and Stanford type A dissections or descending (DTAA) and Stanford type B dissections (Type B). Descending TAAD, also termed Type B dissection can be driven by both syndromic (e.g., hereditary genetic conditions such as familial TAAD, Ehlers-Danlos and Marfan syndromes) and non-syndromic/sporadic (as yet undescribed genetic causes, atherosclerosis, and hypertension)^1^. Syndromic causes of descending TAAD are rare, and a majority of descending TAAD events occur absent any *a priori* indicators of patient risk. While imaging is a highly effective means of detecting and diagnosing TAAD, the low prevalence of TAAD in the general population renders the cost-benefit ratio of such a screening approach prohibitive.

Circulating biomarkers capable of detecting the presence of descending thoracic aneurysm and risk for type B dissection would provide a valuable and cost-effective tool to screen for risk and flag individuals from the general population for more detailed follow up and diagnosis. To date, there are a large proportion of TAAD studies focused on ascending disease, but differences in etiology and other aspects of descending disease warrant focused attention on mechanisms and biomarkers for disease cases in this specific region. Since many times Type B dissections occur absent of predisposing aneurysm formation^5^, determining whether there are distinguishing biomarkers for these unique type B cases will be an important consideration. Dissection-specific biomarkers could also assist in evaluating the progression of descending aneurysms (DTAA) and predicting likelihood for imminent dissection risk.

A number of studies have explored possible biomarkers for thoracic aortic aneurysms and dissections and are the subject of a recent and thorough review^6^. A smaller handful of studies focused specifically on descending thoracic aortic disease^7^. Among markers studied thus far, many have shown preliminary promise including d-dimer, matrix metalloproteinases, certain collagen chains, smooth muscle cell proteins, and various inflammatory markers such including the somewhat general inflammatory marker CRP. These studies have all focused on biomarkers to distinguish aneurysm and/or dissection from normal and/or cases of acute coronary distress not caused by aneurysm or dissection. Biomarkers that can distinguish aneurysm from dissection may also be of clinical interest, as these molecules could aid in therapeutic decisions regarding timing of surgical intervention as aneurysmal tissue progresses toward increasing likelihood for dissection and degeneration. Our recent proteomic analysis of aneurysmal versus dissected descending thoracic aortic tissue found numerous proteins that were differentially expressed between the two groups^8^, however it is unclear whether any of these tissue-derived proteins would be altered in the circulation and indicative of the two disease states.

Machine learning (ML) is a promising tool for automated classification of groups from proteomics data^9^. ML can take any collection of input data and estimate a mathematical function that predicts a categorical outcome, such as the presence or absence of a disease, or a continuous measure like age. Unlike statistical models, which allow for a quantitative measure of confidence for a relationship, ML can find patterns in unwieldy data with nonlinear interactions^10^. ML is also helpful when there are more input variables than number of subjects^10^. Thirdly, ML model interpretation methods can be used to reveal which proteins are relevant for differentiating between two similar diseases^11^. The application of ML to healthcare has enabled discovery of biomarkers associated with cancer, COVID-19 disease severity^12–15^, and subtypes of diseases^16^. ML model interpretation or feature selection methods can be used to reveal which proteins are relevant for model prediction, and prior work suggests that clustering of similar proteins as a feature selection technique before applying ML methods enables accurate disease classification^11^. These successes in the application of ML have led us to explore whether ML models given inputs of plasma proteins could distinguish between aneurysm and dissection.

The goal of the current work was to leverage ML to distinguish the plasma proteomes from two similar diseases not otherwise distinguishable using a standard statistical approach and provide preliminary insight into mass spectrometry detectable, circulating proteomic signatures capable of separating them. Toward this goal, we profiled the plasma proteomes of individuals with DTAA and Type B and applied ML strategies to identify protein features best able to discriminate between them.

## Methods

### Sample Collection and Study Design

Patients with descending thoracic aortic disease were selected retrospectively from a biorepository of aortic disease patients hosted by Dr. Milewicz and team at the University of Texas, Houston. All patients with a well-preserved plasma sample and an isolated diagnosis of DTAA or Type B dissection were selected for proteomic analysis. Blood samples were collected prior to surgery and held in the patient’s room or nurse’s station until transported by the research nurse to our laboratory (in the same day). On receipt, each sample was logged into the computerized biorepository database and labeled with a unique bar-coded identity number and processed into plasma within two hours. The collection tubes were gently inverted 8 – 10 times and then centrifuged at 1650 RCF for 25 minutes at 22°C. The plasma layer of each tube was transferred to labeled 2ml cryovial tubes (0.5mL per tube) and frozen at -80°C until further use. One aliquot per patient was shipped on dry ice to the proteomics research team at Cedars-Sinai Medical Center for plasma proteomic sample preparation and analysis. All patients included in this study provided informed consent and their recruitment and participation was approved by the institutional review boards of UT Houston.

### Sample Preparation for Liquid Chromatography Mass Spectrometry

Proteins from 5uL of plasma were processed for protein denaturation, reduction, alkylation, and tryptic digestion using the manufacturer protocols for the Protifi (Farmingdale, NY) S-Trap protein sample preparation workflow. Resulting peptides were quantified by BCA assay and 2uL of peptide suspension from each sample was pooled to make a master mix used for quality control monitoring purposes and for generation of peptide assay libraries for peptide and protein identification from individual DIA-MS samples (see below).

### Mass Spectrometry Acquisition

#### Individual plasma samples

Mass spectrometry data were acquired on an Orbitrap Exploris 480 (ThermoFisher, Bremen, Germany) instrument with LC separation on an Ultimate 3000 HPLC system using a trap-elute set up on a 150 mm long, 0.3 mm inner diameter reversed phase column (Phenomenex, Luna Polar C18 3 um). A binary analytical gradient using 0.1 % formic acid in water (mobile phase A) and 0.1% formic acid in acetonitrile (mobile phase B) was delivered as follows at 9.5 uL/min: start at 1% B and hold for 2 minutes, ramp to 4%B in 30 seconds, ramp to 12% B over 20 minutes, ramp to 27% B in 24 minutes, ramp to 45% B over 16 minutes (60 minutes total). A separate cleaning equilibration method ran at 98% B for 8 minutes and equilibrated at 2% B for 2 minutes.

The peptides eluted from the analytical column into a Newomics M3 8-nozzle emitter and electrosprayed at 3 kV into a 300°C ion transfer tube temperature. The mass spectrometer was operated in data independent acquisition (DIA) mode acquiring an MS1 scan for 100msec on all ions between 400-1100 m/z and then completing a series of 25msec MS/MS fragment scans on 50 equally spaced 12 m/z width precursor isolation windows. Orbitrap resolution and normalized AGC target were 60,000 and 200% for MS1 and 15,000 and 400% for MS2. Collision energy for HCD fragmentation was set to 30%. The acquisition sequence included repeated sampling of pooled digest to monitor MS QC as well as evenly spaced samples of pooled plasma from across the 3 x 96-well digestion plates to monitor digestion QC.

#### Gas Phase Fractionation based Library Generation

A sample pool was used to generate a spectral library specific to this sample type and analytical platform. Gas phase fractionation limits the scope of the mass spectrometer to a narrow m/z range thus exhaustively fragmenting the corresponding peptide ions and maximizing the probability of their identification and incorporation in the generated library. Multiple injections probing different narrow m/z ranges are compiled to cover the entire range of interest (400 to 1000 m/z). Two complementary approaches were used to generate this library: data dependent acquisition (DDA) of peptides within 120 m/z wide mass ranges (400 to 520 m/z, 520 to 640 m/z … 880 - 1000 m/z) in duplicate; and data independent acquisition (DIA) using 1 m/z wide isolation windows covering 40 m/z at a time (400 to 440 m/z, 440 to 480 m/z … 960 to 1000 m/z). Other than the mass ranges and isolation window widths the method settings were matched to the data acquisition method.

### Proteomic Dataset Generation

#### Library construction

Gas phase fractionated DIA and DDA runs were analyzed using the FragPipe platform^17^. For DIA mode, DIA-Umpire was first used to extract pseudospectra^18^. DIA pseudospectra and DDA spectra were searched separately using the FragPipe workflow to perform spectral matching, PSM probability scoring and then spectral library generation against a Uniprot Human FASTA predicted protein sequence database. Search settings were as follows: mass errors were set to +/-8 ppm for precursor and fragment masses in DDA and +/- 1 m/z for the same masses in the DIA datasets. Carbamidomethylation of cysteine was set as static and methionine oxidation, phosphorylations of serine, threonine, and tyrosine, N-terminal acetylation, pyroQ, pyroC, and pyroE, were selected as variable modifications. Peptides identified at 1% FDR were compiled into a spectral library with EasyPQP. Spectral libraries from the DDA and DIA runs were merged at the level of the final library tsv document. We assumed that a DIA-based peptide identification would provide the best representation of the fragments and their relative intensities for identification within a subsequent DIA run. Thus, only the unique peptides from the DDA library not seen in the DIA runs were appended to the DIA library. The final library contained 8,819 precursors and 407 proteins.

#### Individual subject peptide and protein quantification

Individual DIA runs were processed using DIA-NN^12^ by searching against the sample-specific libraries (described above) using double pass mode and match between runs. Retention time-based normalization setting was used and maxLFQ calculated protein intensities, provided in the main DIANN output matrices, were used for further analysis.

### Data Cleaning

Peptides and proteins with at least one missing or zero value were removed from further analysis, reducing the number of peptides from 8,243 to 1,549 and proteins from 357 to 238 quantified across all samples. Forty proteins with multiple Uniprot identifiers in their group were removed, resulting in 198 proteins measured across all samples. Only patients with complete demographic and sample collection data (i.e., age, sex, race) were carried forward for analysis and four patients were removed who had duplicate rows between both peptide and protein datasets, resulting in 137 patients. Peptide and protein quantities were log_2_ transformed across each sample and corrected for batch effects using pyComBat^19^.

### Statistical Analysis

Data cleaning, analysis, and model training were performed in Python version 3.7.11 (SciKit-Learn^20^, SciPy^21^, seaborn^22^, Matplotlib^23^, Plotly^24^, and Statsmodels^25^). Volcano plots were used to visualize the presence of any differentially expressed proteins and peptides between diseases. Log_2_ fold changes (FC) were calculated by subtracting the log_2_ mean quantity for each protein in the control group from the log_2_ mean quantity in the disease group. P-values were determined using independent two-sample t-tests with Benjamini-Hochberg (BH) multiple hypothesis testing correction. Age was compared between groups using a Wilcoxon Rank Sum Test due to non-normal distribution (Shapiro-Wilks p-value <0.05). Fisher’s Exact Test was used to compare categorical variables (sex and ethnicity) between groups. BH adjusted p-values <0.01 were considered statistically significant. To avoid test data leak into train data when performing feature selection^26^, t-tests were performed only on the 80% train set to select significantly different peptides between groups, and then the prediction was made on the 20% test data filtered for these features.

### Machine Learning and Feature Importance

To account for the presence of correlated proteins within our dataset, we grouped quantitatively similar proteins using hierarchical clustering analysis before performing model interpretation to generate a list of uncorrelated protein groups ranked by their level of importance when classifying between disease states. Using only the 80% train set to avoid biasing the feature selection method with test data^26^, we calculated the Spearman correlation coefficients between each protein pair, converted the correlation matrix to a condensed distance matrix, and applied Ward’s linkage to cluster proteins based on distance. In the dendrogram, the number of vertical lines intersecting a horizontal line drawn at a linkage distance threshold represents the number of clusters at that distance. To identify the linkage distance threshold corresponding to optimal model performance, we trained different ML models with a single, representative protein from each cluster at various distance thresholds. The six supervised ML classification algorithms used were Gradient Boosting Decision Trees (GB)^27^, Support Vector Classification (SVC)^28^, Random Forest (RF)^29^, Extra-Trees (ET)^30^, Logistic Regression (LR)^31^, and K-nearest neighbors (KNN)^32^. Data was split into 80% training and 20% final test sets. To avoid overfitting, the 80% training split was used to tune model hyperparameters via a random search with 5-fold cross validation optimized on F1-score, and the 20% test set was held-out until final evaluation using the best model from the random hyperparameter search. This was repeated with up to 200 random sets of hyperparameters, i.e., up to 1,000 models were trained for each ML method. The best hyperparameters were then used to refit each model with the entire 80% training set before assessing final performance with the 20% test set. The model output was stratified during data splitting to represent the proportion of classes in the whole dataset. Due to imbalanced classes, metrics that focus on the minority class, including precision, recall, F1-score, were chosen to represent generalization performance of the test set. The model with the highest F1-score on the training set and the number of features at the corresponding linkage distance were used for evaluation of the test set. F1-score, PR AUC and accuracy scores using this best model were reported for the test set.

Permutation importance (PI), or a decrease in accuracy score when a single feature’s value is randomly shuffled, was calculated on the test set using the number of features determined at the optimal linkage distance. Each feature’s value was randomly shuffled 10 times and a decrease in accuracy score was calculated each time. The mean of the 10 scores was calculated for each protein and the mean decrease in accuracy scores were ordered from high to low. Proteins with the largest mean decrease in accuracy score were most important to the model’s predictions, and the top 10 proteins were visualized in a box and whisker plot.

### Biological Pathway Analysis

All proteins among the top 10 PI protein groups at selected linkage distance were submitted for pathway analysis. GO Biological Process term enrichment analysis was performed using the ClueGO (version 2.5.9)^33^ application within Cytoscape (version 3.9.1)^34^. GO database release date was 5/25/2022. The default parameters were used, except: GO term fusion was turned on, the threshold for statistical significance was set to <0.0001, and the GO tree interval was set to 3-8. Enriched terms were then manually filtered to keep only non-redundant terms that connected all the proteins to the network.

## Results

**Figure 1A** depicts the study workflow. Plasma samples were obtained from 137 individuals, of which 75 were diagnosed with isolated Type B and 62 were diagnosed with isolated DTAA. Plasma proteomes were generated using DIA-MS and searched against a custom library of sample-specific peptides generated from pooled study plasma samples. Six ML classification algorithms were then trained on the protein quantities from DIA-NN to predict Type B or DTAA disease and feature importance allowed for ranking of most important protein predictors of disease.

**Figure 1.**
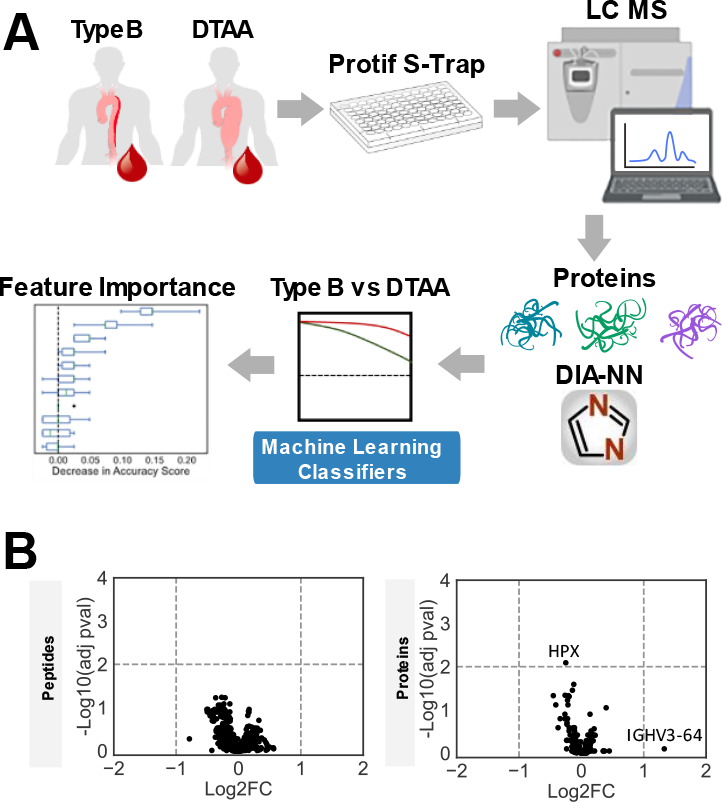
Study overview. **(A)** Proteomic dataset generation and analysis for classification between Type B and DTAA cases and identification of important protein predictors. **(B)** Left: Volcano plot of 1,549 peptides quantified across all samples; no peptides were differentially expressed between diseases. Right: Volcano plot of 198 proteins quantified across all samples; one protein was differentially expressed between diseases. A positive log_2_FC indicates protein mean is higher in Type B. Negative log_10_(adjusted p-value) > 2 corresponds to an adjusted p-value <0.01.

After filtering for the highest quality and most consistent protein identifications, a total of 198 proteins and 1,549 peptides were quantified across all 137 samples included in this study. Volcano plots of the negative log_10_ of the B-H adjusted p-value for all 198 proteins (left) and 1,549 peptides (right) between DTAA and Type B as a function of the log_2_FC of the mean quantities between each group are plotted in **Figure 1B**. A positive fold change indicates the mean protein quantity was higher in Type B relative to DTAA. There were no peptides and only one protein (hemopexin, HPX; p=0.008) that were significantly different between groups. One protein additionally had a log_2_ FC greater than one, immunoglobulin heavy variable-64 (IGHV3-64). Quality control samples were included throughout the data acquisition to account for both digestion and mass spectrometry performance (**Figure S1**).

Patient demographics were compared between groups. Age was significantly higher in the DTAA group while there were no significant differences in sex or ethnicity between Type B and DTAA (**Table 1**). Median age was 57 years in Type B and 66 years in DTAA (p=0.002). Sixty-one percent of patients with Type B and 58% of those with DTAA were male. The most prevalent ethnicity in each group was Caucasian, followed by African American/Black and Hispanic/Latino.

**Table 1.** Baseline characteristics of patients with Type B or DTAA. Continuous variables are reported as median (IQR) and categorical as counts (%).

|  | Type B (N=75) | DTAA (N=62) | p-value |
| --- | --- | --- | --- |
| **Age (yrs)** median (IQR) | 57 (51-67) | 66 (58-74) | 0.002 |
| **Sex** count (%) | M: 46 (61%)<br>F: 29 (39%) | M: 36 (58%)<br>F: 26 (42%) | 0.73 |
| **Ethnicity** count (%) | Caucasian: 37 (49.3%)<br>African American/Black: 24 (32%)<br>Hispanic/Latino: 13 (17.3%)<br>Asian: 1 (1.3%)<br>Other: 0 (0%) | Caucasian: 41 (66.1%)<br>African American/Black: 12 (19.4%)<br>Hispanic/Latino: 7 (11.3%)<br>Asian: 0 (0%)<br>Other: 2 (3.2%) | 0.07 |

When ML models are trained from correlated features, they may learn to rely on one arbitrary representative of the group of correlated features to make predictions. This is especially true of tree-based models. To avoid losing information about the correlated features within a group, a clustering strategy was used before ML model training and interpretation. Using the 80% train set, we visualized correlation between the 198 protein features using a heatmap of Spearman rank-order correlation coefficients. The heatmap indicates distinct groups of highly correlated proteins (**Figure 2A**). Hierarchical clustering of proteins was performed across linkage distances from zero to five; as linkage distance increases, there are fewer uncorrelated protein clusters **(Figure 2B)**. For example, at a linkage distance of four, all proteins are grouped into only two protein clusters. The number of protein clusters at each linkage distance is depicted in **Figure 2C**. A single protein was selected at random from each group of correlated proteins to be used for model training. This allowed tracing each protein selected by the model as ‘important’ back to the larger group of correlated proteins after we performed feature importance on the best model. Average training F1-scores across linkage distances from zero to five were visualized for six ML models **(Figure 2D)**. Training F1-scores generally declined across models as linkage distance increased and there were fewer features input to the models. The highest F1-scoring model using the training set at any linkage distance was SVC with a score of 0.67 at a linkage distance of 0.5. One hundred eleven protein clusters were present at this threshold. **Figure 2E** shows generalizability of the optimized SVC model on the test set (accuracy 0.74, F1-score 0.67, and PR AUC 0.69). Optimal hyperparameters for the SVC model were C=0.1, gamma=1, kernel=poly, probability=True, random_state=42 when tested with the 111 representative proteins from each cluster. PI scores were then calculated for these 111 proteins. **Figure 2F** shows box and whisker plots for the decrease in accuracy score across the 10 permutations for the top 10 most important sentinel proteins sorted by their mean decrease in accuracy score. The mean decrease in accuracy score and representative protein for each cluster are listed in **Table S1**.

**Figure 2.**
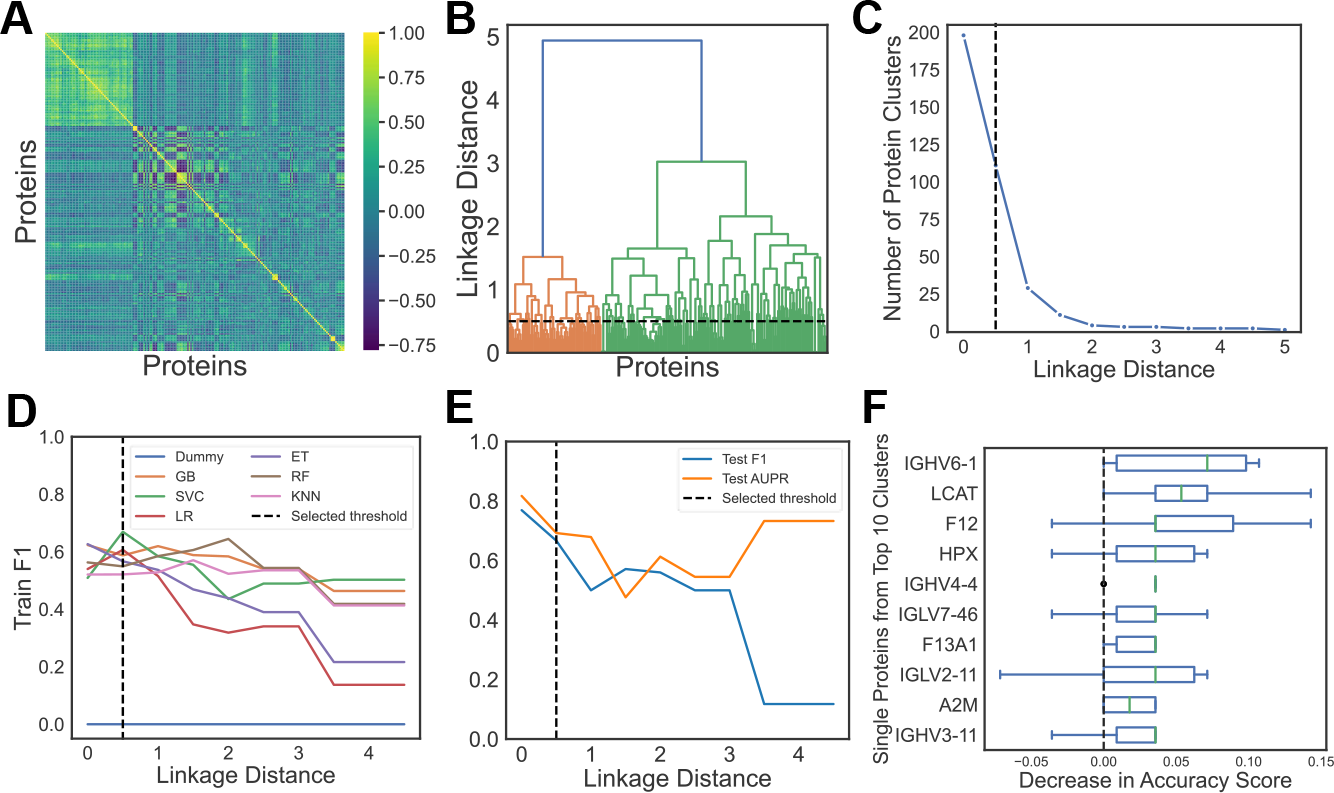
Machine learning (ML) approach for discovery of most predictive proteins between Type B dissection compared to DTAA samples. **(A)** Heatmap of Spearman rank-order correlation coefficients for each pairing between 198 proteins (zero implies no correlation). **(B)** Hierarchical clustering on Spearman rank-order correlations using Ward’s linkage; a linkage distance of 0.5 yields 111 correlated protein clusters. **(C)** Number of uncorrelated protein clusters at each linkage distance. **(D)** Average F1-scores from 5-fold cross validation for each ML model with the training data across linkage distance thresholds from 0 to 5. A single protein from the clusters at each threshold was arbitrarily selected as input to the models; SVC at a linkage distance of 0.5 had the highest F1-score on the train set compared to all other models. At this threshold, there were 111 protein clusters. **E**) F1-score and PR AUC for test set across various thresholds for SVC showing good performance at 0.5 linkage distance threshold (SVC test F1-score 0.67, PR AUC 0.69). (**F)** Box and whisker plots of the distribution of PI scores for the top 10 sentinel proteins from the total 111 clusters. Each box has a line at the median and extends between the lower and upper quartiles of the PI distribution for that protein.

Using these mean PI scores, we filtered for only the proteins with a positive score. The threshold to apply as a cut off to the PI score is objective^10^, i.e., we chose values above zero, but a higher PI may have been selected to filter the number of proteins. There were 23 sentinel proteins with a positive PI score and a total of 38 proteins making up the 23 sentinel protein clusters. PI scores provide a ranked list of important proteins in prediction between the two diseases, versus statistics which selects a protein based on adjusted p-value cut-off.^10^ To visualize the difference in level of information provided by each method, negative log_10_ of the adjusted p-values derived for each protein between the two groups was plotted as a function of the mean PI scores for these 23 proteins (**Figure 3**). While HPX is the only significantly different protein, there are proteins (LCAT, F12 and IGHV6-1) chosen by the model as more ‘important’ than for classification between diseases. There are also a number of proteins that have similar mean PI scores to HPX. Log_2_FC, B-H adjusted p-values and mean PI scores for the top 10 protein clusters, mapping to 19 total proteins, depicted in **Figure 2F** are listed in **Table 2**. IGHV6-1 had the highest score, however it’s quantity between groups was not significantly different (B-H adjusted p-value 0.38). A log_2_FC of -0.17 indicates the mean quantity of IGHV6-1 was higher in the DTAA group compared to the Type B group. Mean quantities per group, log_2_FC, negative log_10_ adjusted p-values and adjusted p-values for all 198 proteins are listed in **Table S2**.

**Table 2.** Log_2_FC, adjusted p-values, and mean PI scores for 10 highest PI scoring clusters. Representative cluster protein is shown in bold for each cluster. Positive log_2_FC indicates mean protein quantity is higher in Type B group. Negative log_2_FC indicates mean protein quantity is higher in DTAA group.

| Clustered Proteins | Log<sub>2</sub>FC | B-H adjusted p-value | Mean PI |
| --- | --- | --- | --- |
| **IGHV6-1** | -0.17 | 0.38 | 0.057 |
| **LCAT**<br>HRG<br>HGFAC<br>PGLYRP2 | -0.21 | 0.047 | 0.057 |
| **F12** | -0.21 | 0.076 | 0.050 |
| **HPX** | -0.25 | 0.0081 | 0.032 |
| **IGHV4-4**<br>IGHV1-45<br>IGHV1-24<br>IGHV5-51<br>IGLV1-40 | -0.055 | 0.89 | 0.029 |
| **IGLV7-46** | -0.070 | 0.95 | 0.025 |
| **F13A1**<br>F13B | -0.24 | 0.50 | 0.025 |
| **IGLV2-11** | 0.071 | 0.89 | 0.021 |
| **A2M**<br>FBLN1 | -0.14 | 0.55 | 0.018 |
| **IGHV3-11** | -0.10 | 0.84 | 0.018 |

**Figure 3.**
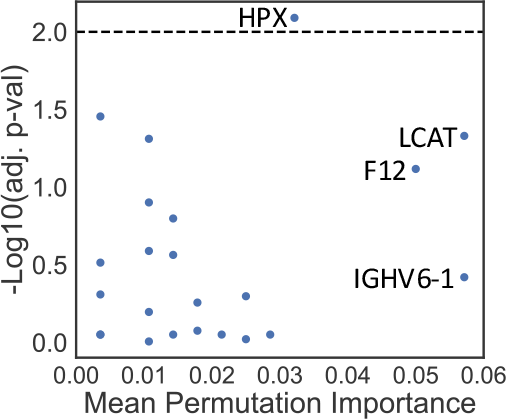
Comparison of -log10 (adjusted p-values) to mean PI for all proteins with a positive mean PI score.

These 19 proteins from the top 10 clusters shown in **Figure 2F** were then input to GO term enrichment analysis. Proteins in the following pathways were significantly enriched in the plasma of Type B dissection compared to DTAA patients: complement activation, humoral immune response mediated by circulating immunoglobin, and blood coagulation/fibrin clot formation (**Figure 4)**.

**Figure 4.**
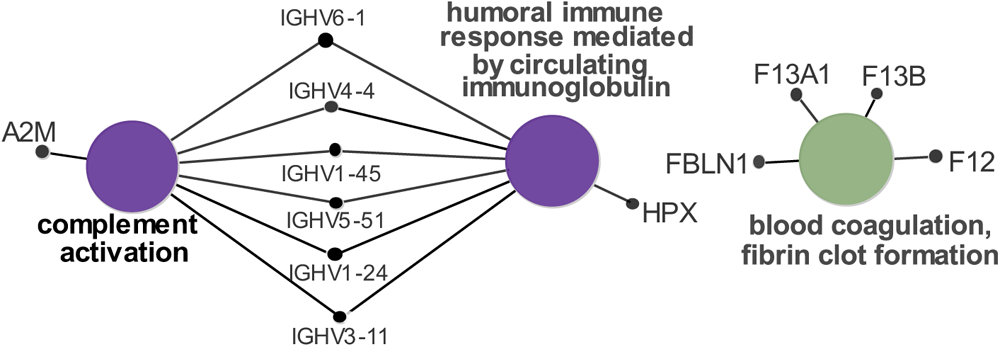
Biological term enrichment analysis of proteins altered between Type B dissection vs DTAA samples.

## Discussion

These data represent the most comprehensive analysis we are aware of describing the circulating proteome from patients with descending aortic disease to date. As the process of identifying informative features for eventual biomarker panel production is arguably more intuitive at the protein level and proteins appeared highly correlated, we segregated proteins into similar clusters, and used permutation-based importance ranking to identify correlated protein groups that were informative for separating patients between DTAA and Type B conditions. We were able to find only one differentially expressed protein between Type B and DTAA patients, yet the ML approaches still differentiated patients between these two groups to some extent (AUPR=0.7) using the test set data not used during model training. We found that we can reduce our protein data to a list of 111 uncorrelated proteins to train the highest performing model. Clusters important in distinguishing between the diseases included proteins involved in inflammation and coagulation.

ML is likely picking up patterns across many measured proteins, compared to statistical tests that ask if one protein in aggregate has a different mean value between the groups. This may be useful for complex diseases that are heterogeneous across individuals; ML models can learn multiple signatures leading to disease. The main downside of using ML is that it requires many samples, typically hundreds, compared to statistics, which can be performed with as few as three replicates per group. Thus, modeling larger proteomic datasets using more sophisticated and modern approaches may be a potent approach for gaining new insight into the power of proteomic signatures for predictive biomarker development.

Many of the informative proteins selected by the ML model demonstrated similar trends for differential abundance in our previous proteomic analysis of tissue samples comparing Type B and DTAA ^8^. Plasma HPX was both a ML model selected and significantly abundant protein between aneurysm and dissection cases in our study, and also demonstrated a trend toward increased abundance in aneurysm tissue relative to dissected tissue. Hemopexin is a heme scavenging protein considered to be generally protective against cardiovascular disease and atherosclerosis^35,36^. Similarly matched trends for abundance in both circulating plasma and tissue proteome of DTAA relative to Type B patients was observed for another heme scavenger, A2M, as well as proteins IGHV6-1, HRG, PGLYRP2 and F13A and B. Prominent involvement of immunoglobulins including IGHV6-1 is consistent with recent reports of a potentially pathogenic role for B cells and immunoglobulin deposition in abdominal aortic aneurysm (AAA)^37^, and suggests similar involvement in the descending thoracic aorta. Factor 13A and B are fibrinolytic proteins with gene polymorphisms associated with AAA and hemolytic aneurysmal subarachnoid hemorrhage in the brain^38,39^ . One other interesting standouts in the list of informative proteins differentiating aneurysm and dissection was SAA4 (elevated in dissection). Overall levels of circulating Serum Amyloid A were recently identified as a potential biomarker for acute ascending and type B aortic dissection^40^. While the prior mentioned study did not differentiate between SAA subtypes (e.g., SAA1, SAA2, or SAA4), this work generally supports the biological relevance of SAA4 protein as potentially important for distinguishing aneurysm from dissection in descending thoracic aortic disease. Taken together, many of the proteins selected by the ML models as highly informative for discriminating diseases are supported by solid corroborating biological evidence and for some, prior identification as putative biomarkers for thoracic aortic disease, thus providing evidence for the validity of this approach for identifying informative plasma biomarker candidates.

This study is a preliminary effort to address a pressing need for informative biomarkers for descending thoracic disease, and while powerful and biologically plausible new hypotheses have been generated, there are some weaknesses to mention. It is likely that small sample sizes impacted discriminative power and performance of the ML classifier, and future studies that expand the numbers of patients are needed. Samples were collected at very late-stage disease, just prior to surgical intervention. By this time, many aneurysm patients may have very similar overall pro-inflammatory plasma proteome signatures relative to aortic dissection patients. While this can be helpful in distinguishing disease states at their most extreme, the highest translational and clinical impact will come from biomarkers that can detect and distinguish disease at very early stages of development and thus both predict adverse progression and provide theranostics to monitor effectiveness of pharmacological intervention. In addition, dissection absent a prior aneurysm may represent a very distinct pathogenic process for which late-stage aneurysm biomarkers cannot predict, and from which biomarkers of Type B dissection alone will not transfer to cases of dissection after significant aneurysm degeneration. Thus, future work is needed to determine the robustness of the selected candidate markers in additional patients at later disease stage and, importantly, then determine which putative biomarkers may be informative at detecting early-stage disease and predicting risk for severe outcomes.

## Conclusions

The data presented in this preliminary report provide a framework and preliminary protein signature from which ongoing efforts will be built and support the power of ML for identifying biomarker candidates and building discriminative models to distinguish between biological states within the context of descending thoracic aortic disease.

## Supplemental Figure Legend

**Figure S1**. Quality Control analysis of digestion and mass spectrometry reference plasma pools

## Supplemental Table Legends

**Table S1**: 111 representative cluster proteins and all the correlated proteins within each cluster; the 111 representative proteins were used as inputs to the highest performing model classifying between disease. (xlsx)

**Table S2**: Mean protein quantities in each disease group, mean difference, log_2_(Type B/DTAA), negative log_10_ adjusted p-values and adjusted p-values for all 198 proteins. Positive log_2_(Type B/DTAA) indicates mean is higher in Type B group. Negative log10(adjusted p-value) > 2 corresponds to an adjusted p-value <0.01. (xlsx)

## Declarations

### Ethics approval and consent to participate

All patients included in this study provided informed consent and their recruitment and participation was approved by the institutional review boards of UT Houston.

### Consent for publication

Not applicable.

## Availability of data and materials

The mass spectrometry proteomics data have been deposited to the ProteomeXchange archive via the PRIDE partner repository with the data set identifier PXD041337 and 10.6019/PXD041337.

Username: The password for the data will be made public upon manuscript acceptance.

The python code for data analysis in JupyterLab notebooks can be found at: https://github.com/xomicsdatascience/Aneurysm-ML.

## Competing interests

The authors declare that they have no competing interests.

## Author contributions

Conceptualization, A.M., J.G.M., S.P., A.A., D.M., D.G; Methodology, A.M., J.G.M., S.P., S.K., M.A., D.G.; Software, A.M., J.G.M.; Formal Analysis, A.M., J.G.M., S.P.; Investigation, A.M., J.G.M., S.P., M.A., S.K.; Resources, J.G.M., S.P., Data Curation, A.M., J.G.M., S.P.; Writing - Original Draft. A.M., J.G.M., S.P.; Writing - Reviewing & Editing, A.M., J.G.M., S.P., S.K., M.A., D.G., D.B., K-Y.C., P.K.S., D.M., A.A.; Visualization, A.M., J.G.M., S.P.; Supervision, J.G.M., S.P.; Project Administration, A.M., J.G.M., S.P., A.A., D.M., D.B.; Funding Acquisition, J.G.M., S.P., A.A., D.M.

## Funding

This work was supported by the Cedars-Sinai Leon Fine award for Translational Research (*S.P, A.A)* as well as grants from the NIH NIGMS (*R35 GM142502 J.G.M.)*, the NIH NHLBI *(R00HL128787, S.P.)* and the NIH NHLBI *(1P01HL110869, D.M.)*.

## Acknowledgements

The authors thank Dasom Hwang for help with graphic design and the Cedars-Sinai Proteomics and Metabolomics Core facility for support with Mass Spectrometry data acquisition.

## Notes

### Competing Interest Statement

The authors have declared no competing interest.

### Summary of Updates

The first version of the manuscript compared our disease groups with data from healthy people that was collected at a different site. Differences in sample collection likely resulted in systematic technical differences between the proteomic data from the healthy group relative to the disease groups. The revised version removes this group of healthy people, and instead focuses on the comparison between the disease groups that were sampled in the same way.

