## supplemental figure 1 for "Differentiation between Descending Thoracic Aortic Diseases using Machine Learning and Plasma Proteomic Signatures"

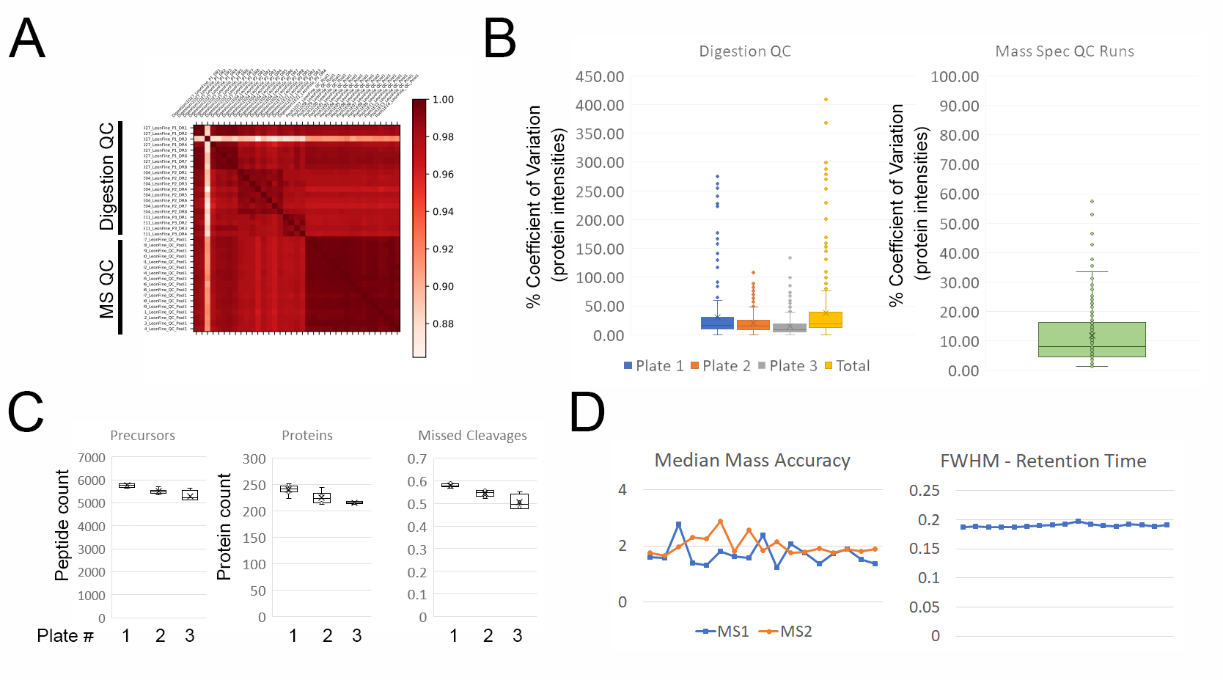

**Supplemental Figure 1**. Quality control analysis of reference plasma pools for monitoring digestion and mass spectrometry performance. **A.** Sample correlations between separate digestions of the same control plasma pool spread throughout all processing plates (digestion QC) and separate mass spectrometry runs on the same pooled digest (MS QC) spread throughout the total acquisition period. **B.** Coefficient of Variation (%CV) for QC samples within and across processing plates (left panel) and MS QC runs (right panel). **C.** Protein and peptide identifications across the processing plates. **D.** Mass accuracy and chromatography performance (full width half max) across all MS QC runs.
